# Isolation and characterization of antagonistic bacteria with the potential for biocontrol of soil-borne wheat diseases

**DOI:** 10.1101/325647

**Authors:** Xiaohui Wang, Changdong Wang, Chao Ji, Qian Li, Jiamiao Zhang, Xin Song, Sui Jun Kang, Zhaoyang Liu, Jianfeng Du, Xunli Liu

## Abstract

*Bacillus amyloliquefaciens* subsp. *plantarum* XH-9 is a plant-beneficial rhizobacterium that shows good antagonistic potential against phytopathogens by releasing diffusible and volatile antibiotics, and secreting hydrolytic enzymes. Furthermore, the XH-9 strain possesses important plant growth-promoting characteristics, including nitrogen fixation (7.92 ± 1.05 mg/g), phosphate solubilization (58.67 ± 4.20 μg/L), potassium solubilization (10.07 ± 1.26 μg/mL), and the presence of siderophores (4.92 ± 0.46 μg/mL), indole-3-acetic acid (IAA) (7.76 ± 0.51 μg/mL) and 1-aminocyclopropane-1-carboxylic acid deaminase (ACC-deaminase) (4.67 ± 1.21 nmol/[mg•h]). Moreover, the XH-9 strain showed good capacities for wheat, corn, and chili root colonization, which are critical prerequisites for controlling soil-borne diseases as a bio-control agent. Real-time quantitative polymerase chain reaction experiments showed that the amount of *Fusarium oxysporum* DNA associated with the XH-9 strain after treatment significantly decreased compared with control group. Accordingly, wheat plants inoculated with the XH-9 strain showed significant increases in the plant shoot heights (14.20%), root lengths (32.25%), dry biomass levels (11.93%), and fresh biomass levels (16.28%) relative to the un-inoculated plants. The results obtained in this study suggest that the XH-9 strain has potential as plant-growth promoter and biocontrol agent when applied in local arable land to prevent damage caused by *F. oxysporum* and other phytopathogens.

**Importance:** Plant diseases, particularly soilborne pathogens, play a significant role in the destruction of agricultural resources. Although these diseases can be controlled to some extent with crop and fungicides, while these measures increase the cost of production, promote resistance, and lead to environmental contamination, so they are being phased out. Plant growth-promoting rhizobacteria are an alternative to chemical pesticides that can play a key role in crop production by means of siderophore and indole-3-acetic acid production, antagonism to soilborne root pathogens, phosphate and potassium solubilization, and nitrogen fixation. These rhizobacteria can also promote a beneficial change in the microorganism community by significantly reducing its pathogenic fungi component. Their use is fully in accord with the principles of sustainability.

## Introduction

Plant diseases induced by soil-borne plant pathogens are among the most difficult to control and create challenging problems in agriculture. The *Fusarium* genus comprises several fungal species, many of which may cause crown rot, stem rot, and head blight in wheat (1). Especially, *Fusarium oxysporum* causes vascular wilt and root rot in over 100 plant species (2). In the past few decades, measures for preventing and controlling soil-borne diseases have usually relied on the use of resistant varieties and fungicides. However, developing resistant varieties can be difficult in the absence of a dominant gene (3), and chemical fungicides represent a potential risk to both the environment and human health (4). Furthermore, previous reports have shown that chemical-control measures create imbalances in the microbial community, which may be reduce the activity of beneficial organisms and may also lead to the development of resistant pathogen strains (3). In this regard, biological control by using plant growth-promoting rhizobacteria (PGPR) is a potentially viable alternative strategy for disease management (5).

PGPR have been reported to improve plant growth and suppress the growth of plant pathogens (6). Several mechanisms have been suggested whereby bacteria can promote plant growth (phytohormone production, nitrogen fixation, and stimulating nutrient uptake) and reduce plant diseases (competition for space, nutrients, and ecological niches; production of hydrocyanic acid, siderophores, and antifungal compounds; or the induction of systemic resistance [ISR] (7, 8). Furthermore, the ability to colonize the rhizosphere has been considered a relevant trait for biocontrol (9).

To date, several microorganisms have been successfully used to control plant pathogens, of which *Pseudomonas* spp. and *Bacillus* spp. are important they are aggressive colonizers of the rhizosphere of various crops and show a broad spectrum of antagonistic activities against many pathogens (10). In addition, nonpathogenic bacteria, fungi, and actinomycetes belonging to the genera *Trichoderma, Fusarium, Streptomyces*, and *Serratia* have been used as potential biocontrol agents against damping-off and root rot pathogens, and other soil-borne pathogens (11–13).

Therefore, the aims of this study were to: (1) isolate and evaluate indigenous soil bacteria for their biocontrol potentials against fungal pathogens; (2) identify and characterize select antagonistic bacteria based on the 16S rDNA gene sequences; (3) reveal the mechanism for promoting plant growth and suppressing plant pathogens in vitro; (4) test the ability of bacteria to colonize the rhizosphereof different crops; and (5) determine the impact of selected strain on the *F. oxysporum* content in the wheat rhizosphere by performing realtime quantitative polymerase chain reaction (qPCR) experiments.

## Results

### Isolation, characterization, and identification of antagonistic bacteria

Numerous bacterial colonies with obviously variable morphological features were grown on Potato dextrose agar (PDA) medium. Several isolates of effective bacterial
antagonists belonged to the *Bacillus* genus. Subsequently, these bacterial isolates were evaluated for their plant growth-promoting traits. Among them, the XH-9 strain was selected from our collection in this work, which not only showed wide broad-spectrum activity against phytopathogens (Fig. 1), but also showed evidence of nitrogen fixation (7.92 ± 1.05 mg/g); phosphate solubilization (58.67 ± 4.20 μg/L) and potassium solubilization (10.07 ± 1.26 μg/mL); and siderophore (4.92 ± 0.46 μg/mL), IAA (7.76 ± 0.51 μg/mL), and ACC deaminase (4.67 ± 1.21 nmol/[mg•h]) production. Thus, the XH-9 strain possessed several key plant growth-promoting characteristics. Next, the XH-9 strain was identified as *Bacillus amyloliquefaciens subsp. Plantarum*, based on phylogenetic analysis of the 16S rRNA sequence and was deposited in the China General Microbiological Culture Collection Center (CGMCC) under accession number no.13151.

**Fig. 1.**
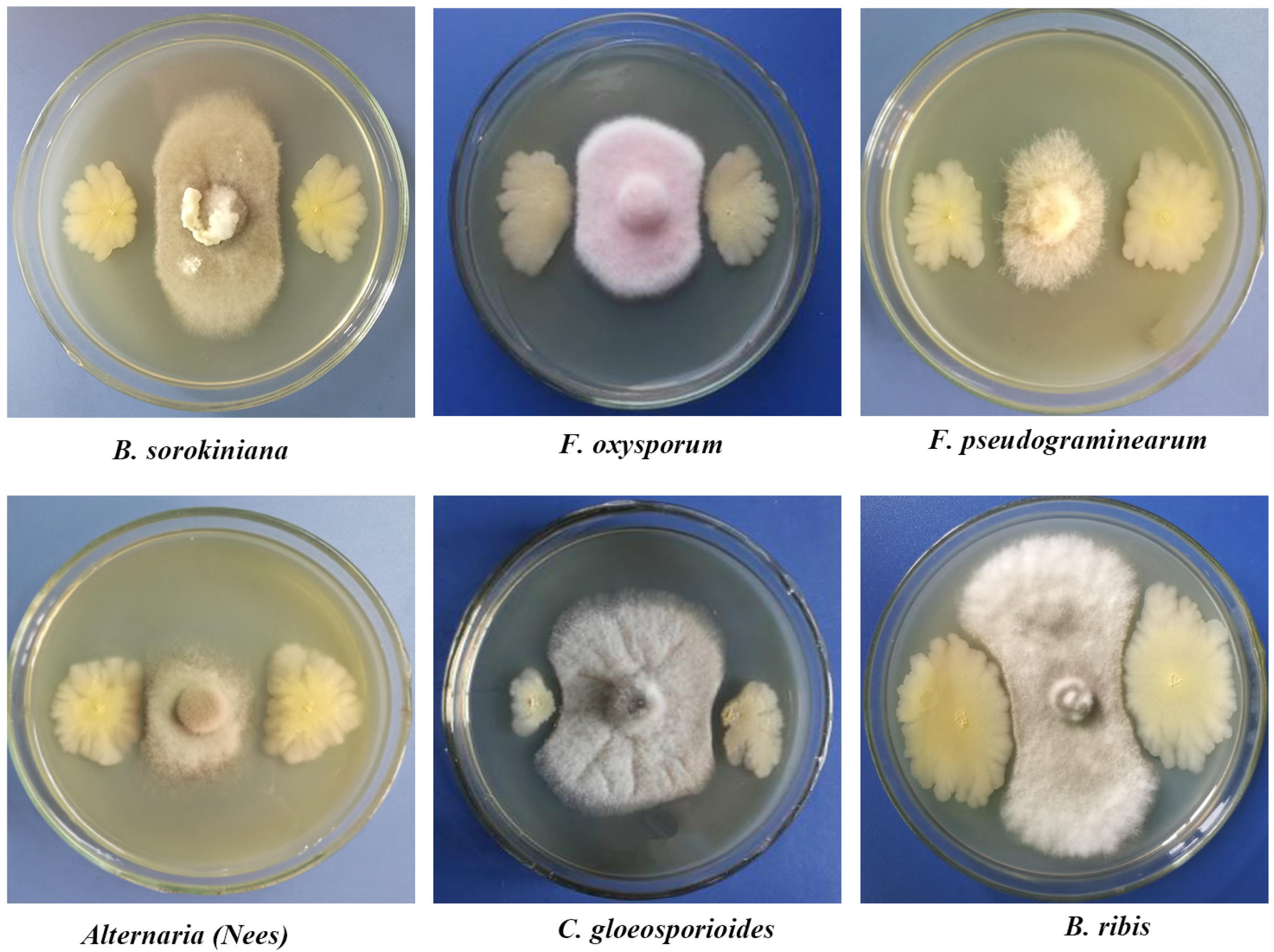
In vitro antagonistic activity of *Bacillus amyloliquefaciens* subsp. *plantarum* (XH-9 strain) against selected pathogen in dual culture assays on PDA medium, 4 d after incubation.

### Antagonistic mechanisms

Both diffusible antibiotics and volatile metabolites of the XH-9 strain inhibited the growth of *F. oxysporum, B. sorokiniana*, and *R. solani.* The colony diameters of these pathogens in the treatment and control groups are presented in Table 1. Compared with the control group, diffusible antibiotics and volatile metabolites efficiently reduced the growth of *F. oxysporum* to 6.59% and 47.07%, respectively, and the grown of *R. solani* to 14.29% and 54.29%, respectively. Both diffusible antibiotics and volatile metabolites completely inhibited the growth of *B. sorokiniana*. Additionally, the XH-9 strain grew in solid media containing sodium carboxymethyl cellulose, yeast glucans, or skimmed milk as the sole carbon source and produced hyaline zones around the bacterial colonies (Fig. 2). These data indicated that the XH-9 strain produced several hydrolases. Quantification of the produced enzymes showed that the cellulase, glucanase, and protease levels reached 22.42 ± 2.63 U/mL, 624.19 ± 52.42 U/mL, and 13.40 ±1.28 U/mL, respectively. Interestingly, the level of chitinase secreted by the XH-9 strain was up to 769.21 ± 27.90 U/mL, although no semicircular hyaline zone formed around the colonies on the plate containing colloidal chitin.

**Fig.2.**
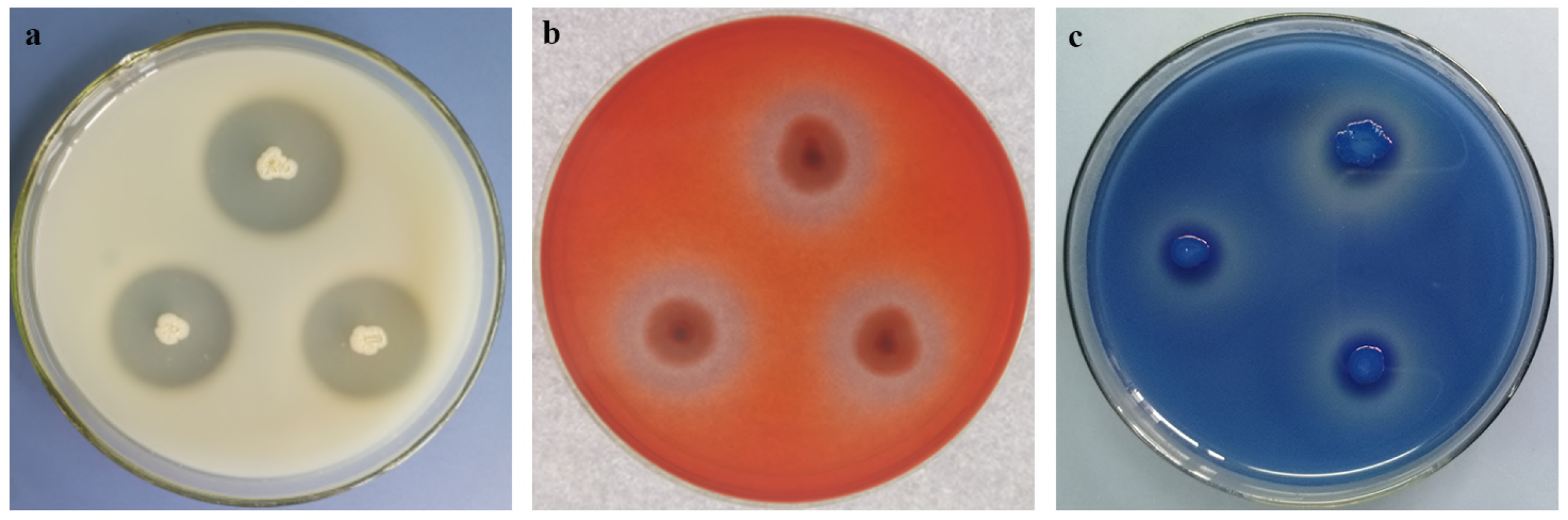
Fungal cell wall-degrading enzymes produced by XH-9 strain. (a) Protease activity analyzed using skim milk agar plates. (b) Cellulase activity analyzed using carboxyl methyl cellulose agar plates. (c) Glucanase activity analyzed using Pachyman solid medium supplemented with 6% aniline blue.

**Table 1.** Diffusible and volatile antibiotics secreted by XH-9 strain against phytopathogens

| Treatment | Diameter of the colonies (cm) |  |  |
| --- | --- | --- | --- |
|  | *Fusarium oxysporum* | *Rhizoctonia solani* | *Bipolaris sorokiniana* |
| CK | 4.1±0.00a | 7.0±0.15a | 3.5±0.00a |
| Diffusible antibiotics | 3.83±0.05b | 6.0±0.06b | 0.50±0.00b |
| Volatile antibiotics | 2.17±0.05c | 3.20±0.10c | 0.50±0.00b |

Note: Values are mean values ± standard deviations (n=3), Means sharing a common letter within the same column are not significantly different at $P < 0.05$ .

### Root-colonization assay

Pot experiments were performed to test the colonization capability of the XH-9 strain. We assessed bacterial population densities in the rhizosphere of different crops at 10-d intervals after bacterial inoculation. The results shown in Table 2 indicated that the XH-9 strain could colonize chili corn and whea. Almost 10^4^-10^5^ (CFUs)/g of root were detected in root rhizosphere soil samples from different crops. The result indicated that the XH-9 strain had an exceptionally high ability to stably colonize the root system of different crops, which is a critical prerequisite for the successful use of PGPR.

**Table 2.** Population density of XH-9 strain on the different crops rhizosphere (10^5^cfu.g^−1^ dry weight of soil)

| Species | Days after application (d) |  |  |  |  |
| --- | --- | --- | --- | --- | --- |
|  | 10d | 20d | 30d | 40d | 50d |
| wheat | 5.70±0.5a | 4.08±0.32b | 0.61±0.06c | 0.44±0.06c | 0.63±0.03c |
| corn | 3.22±0.22ab | 3.53±0.37a | 2.89±0.51b | 0.80±0.10c | 1.02±0.14c |
| chili | 4.63±0.37a | 0.29±0.03b | 0.30±0.09b | 0.27±0.07b | 0.32±0.04b |

Note: Values are mean values ± standard deviations (n=15), Means sharing a common letter within the same column are not significantly different at $P < 0.05$ .

### Biomass accumulation in a pot experiment

The performance of the XH-9 strain was evaluated after challenging it by inoculation with a suspension of 10^7^ conidia mL^−1^ from *F. oxysporum.* As shown in Table 3, the XH-9 strain promoted the growth of wheat seedlings. Compared with the control group, the XH-9 strain significantly increased wheat growth in terms of the plant shoot height (14.20%), root length (32.25%), dry biomass (11.93%), and fresh biomass (16.28%), relative to the pathogen control.

**Table 3.** Effect of the XH-9 strain on growth parameters of wheat seedling

| Treatment | Shoot height (cm) | Root length (cm) | Fresh weight (g) | Dry weight (g) | Root/shoot ratio |
| --- | --- | --- | --- | --- | --- |
| CK | 16.2±0.94b | 9.83±0.60b | 3.01±0.15b | 0.503±0.06 b | 0.352±0.01b |
| XH-9 | 18.5±0.68a | 13.0±1.69a | 3.50±0.34a | 0.563±0.63a | 0.392±0.011a |

Notes: Values are means ± SD (n = 15). Means sharing a common letter within the same column are not significantly different at $P < 0.05$ . “XH-9” denotes wheat seedlings treated with a suspension of strainXH-9 cells. “CK” denotes wheat seedlings treated with an equal volume of sterile water.

### Plasmid construction and extraction

Genomic DNA was extracted from *F. oxysporum, F. pseudograminearum, F. graminearum, B. sorokiniana*, and *R. solani* and observed under ultraviolet (UV) light and after running it on a 2% agarose gel (Fig. 3a). Next, the JB-F and JB-R primer pair was tested in amplification reactions against all prepared fungal genome DNA samples and water (as a negative control). As expected, the primers only amplifying the target sequence of *F. oxysporum* with a specific PCR product of 323 base pairs (bp) (Fig. 3b), which was used to construct recombinant plasmid. Subsequently, the plasmids were transformed into phage resistant, chemically competent Trans1-T1 cells and grown in PDA medium containing 8 μL isopropyl β-D-1-thiogalac topyranoside (IPTG) and 40 μL X-gal at 30°C. After 24 h, a large number of white and blue colonieals appeared on the plate (Fig. 3c)everal white colonies were selected and sequenced at a commercial sequencing company (Sangon Biotech, Shanghai, China) using the M13 forward and M13 reverse primer. The specific sequence of *F. oxysporum* is shown in additional file. Transformants that were successfully identified were expanded in culture to extract plasmids (TransGen Biotech, Beijing, China), which were used in the standard curve.

**Fig. 3.**
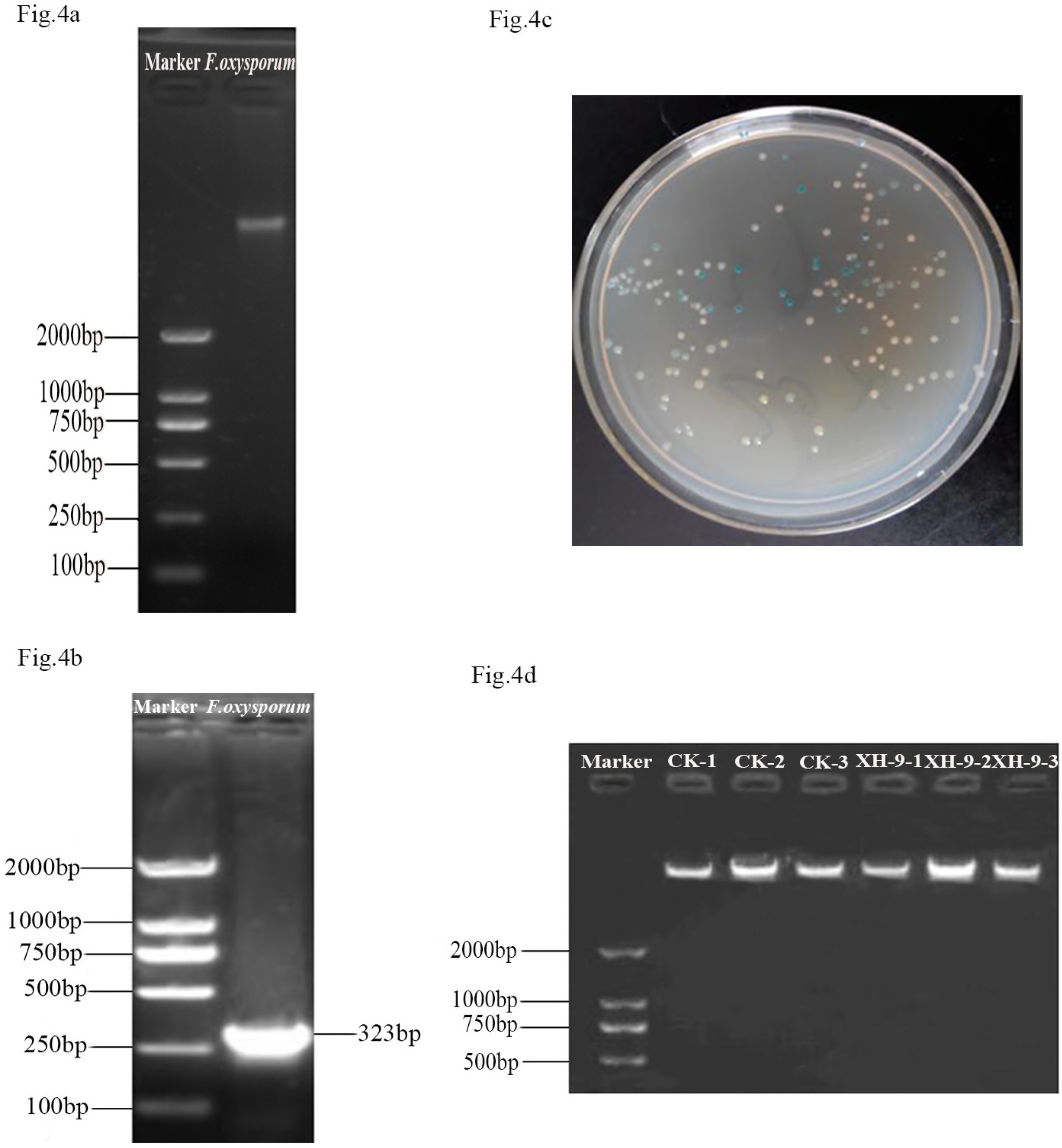
Real-time PCR for quantification of *F. oxysporum*. (a) Fungal genomic DNA was extracted and observed under UV light by running it on a 2% agarose gel. Marker: Trans2K TM Plus II DNA. Lanes 1-5 represent *F. oxysporum, F. pseudograminearum, F. graminearum, B. sorokiniana*, and *R solani*, respectively. (b) A specific fragment of *F. oxysporum* was amplified using the JB-F/R primer pair, which generated a 323-bp DNA product. Marker: Trans2K TM Plus II DNA. Lane 1: sterile water (negative control). Lanes 2-6: *F. oxysporum, F.pseudograminearum, F. graminearum, B. sorokiniana* and *R. solani* respectively. (c) Traditional blue/white screening technique used tothe rapidly and conveniently detects recombinant plasmids. d) Total soil DNA of after control and XH-9 strain treatment. Lanes 1- 3 represent triplicate control group samples and lanes 4-6 represent triplicate XH-9 strain-treated samples. Marker: Trans2K TM Plus II DNA.

### Real-time PCR for the quantification of F. oxysporum

A standard curve was established by plotting the log of known concentrations of *F. oxysporum* DNA against the associated Ct values, with an R^2^ value higher than 0.99 (Fig. 4b). Fig. 4a shows a melting curve of the amplification products, which indicated the specificity of the amplification product. For the rhizosphere soil samples, total soil DNA was extracted (Fig. 3d) and used as a template in qPCR assays. The Ct values observed with unknown soil samples were interpolated with the standard curve, and the calculated *F. oxysporum* DNA concentration from the soil samples was record in Table 4. Compared with the control group, the XH-9 strain significantly inhibited the growth and reproduction of *F. oxysporum* in the wheat rhizosphere at definable intervals. In the control group, the amount of *F. oxysporum* drastically decreased to 3.74 (±0.26) ×104 CFUs/g of soils during the first 20 d, and thereafter, it gradually increased and reached 46.42 (±1.96) × 10^4^ CFUs/g of soil after 50 d. Following treatment with the XH-9 strain, the amount of *F. oxysporum* decreased drastically, being gradually reduced during the first 20 d, and then gradually declining and stabilizing until the final concentration (1-2 × 10^4^ CFUs/g of soil) was relatively stable.

**Table 4.** Contents of Fusarium oxysporum in wheat rhizosphere (×10^4^ cfu.g^−1^ dry weight of soil)

| Days after inoculation XH-9 strain (d) |  |  |  |  |  |
| --- | --- | --- | --- | --- | --- |
| Treatment | 10d | 20d | 30d | 40d | 50d |
| CK | 74.81±4.67a | 3.74±0.26a | 38.46±3.15ba | 39.12±2.085a | 46.42±1.96a |
| XH-9 | 37.01±1.93c | 1.21±0.61c | 0.9±0.06c | 1.08±0.13 b | 1.51±0.58b |

Notes: Values are means ± SD (n = 15). Means sharing a common letter within the same column are not significantly different at $P < 0.05$ . “XH-9” denotes wheat seedlings treated with a suspension of strainXH-9 cells. “CK” denotes wheat seedlings treated with an equal volume of sterile water.

**Fig. 4.**
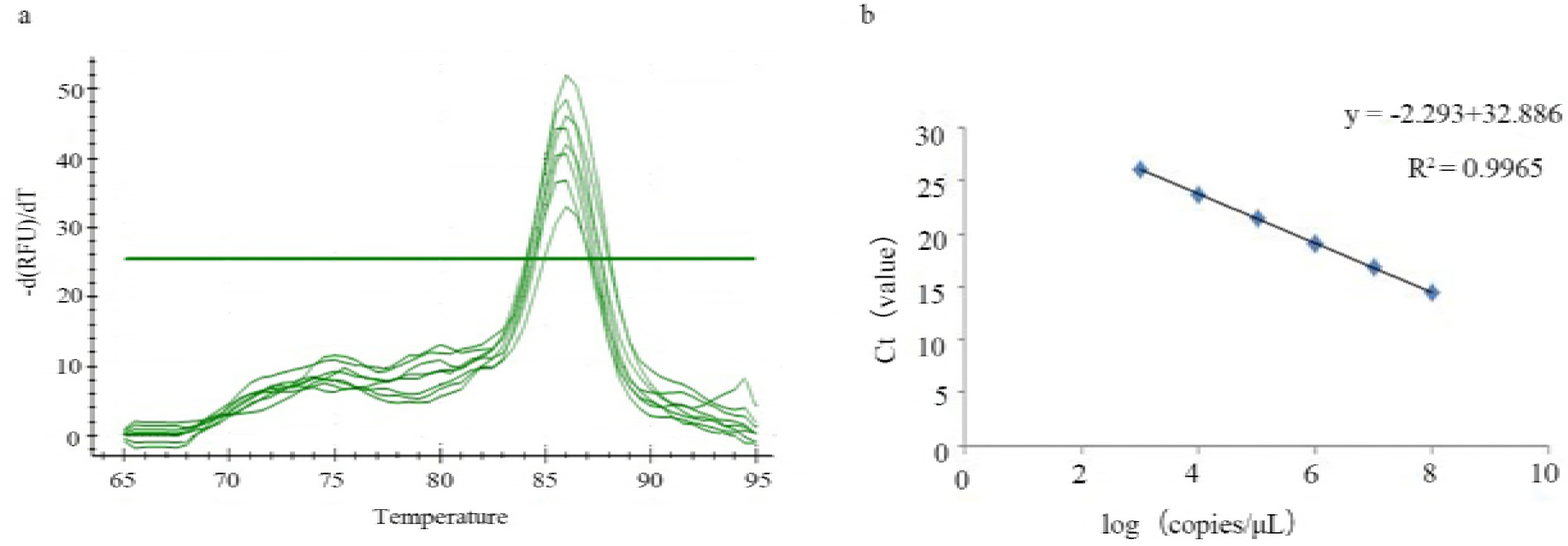
Real-time PCR quantification of *F. oxysporum* in the wheat rhizosphere following XH-9 treatment and control treatment. (a) Melting curve analysis of PCR products from *F. oxysporum.* (b) qPCR standard curve

## Discussion

In the absence of targeted plant-disease control approaches, soil-borne pathogens continue lead to significant crop losses worldwide in both conventional and organic field production systems (11). Development of biological control measures using antagonistic bacteria is one of the most attractive alternatives to chemical fungicides, although it has many limitations including environmental concerns, food quality issues, and tolerance to antifungal agents after long-term applications (14). A trend exists in terms of utilizing microorganisms isolated from rhizosphere soil as biological agents, which may be better adapted to colonize the rhizosphere of crop plants and show better disease control than organisms isolated from the other sources, such as seawater, compost, and silt (15, 16).

Several species of nonpathogenic rhizobacteria, mainly belonging to the *Pseudomonas, Bacillus*, and *Paenibacillus* genera, have been identified from soils and used as potential biocontrol agents against soil-borne pathogens. For example, *B. amyloliquefaciens* FZB45 has been found to be an effective biocontrol agent with P-mobilizing capabilities (17). Treating seeds with *Pseudomonas fluorescens* enhanced resistance to tomato damping off (18). *Paenibacillus polymyxa* E681, with its plant-growth promotion and root-colonization ability, showed promise as aing biocontrol agent for cucumber and barley (19). However, *Bacillus* species have attracted more attention than other species because their long-term viability facilitates the development of commercial products (20). Previous data suggest that *Bacillus* species exert antagonistic activities against pathogens by synthesizing antimicrobial peptides, secreting lytic enzymes, competing for nutrients and space, and inducing systemic resistance (21).

At present, *B. amyloliquefaciens*, which is known promote plant growth and to produce diverse secondary metabolites (22), was studied widely and applied as a biofertilizer and biocontrol agent in agriculture. Compared with other representatives of endospore-forming *B. amyloliquefaciens, B. amyloliquefaciens* subsp. *plantarum* was distinguished by the ability to efficiently colonize plant rhizospheres and su ppress competing phytopathogenic bacteria and fungi (20). For example, *B.amyloliquefaciens plantarum* GR53 mitigated biotic stress by regulating the levels of plant stress hormones and antioxidant activity (21). In addition, the FZB42 strain of *B. amyloliquefaciens* subsp. *Plantarum* was used to enhancing the yield of tomatoes, based on positive results for producing indole compounds and siderophores, and for their biocontrol functions against plant pathogens (23, 24). The S499 strain of *B. amyloliquefaciens* subsp. *Plantarum* was applied as a plant-beneficial rhizobacterium with good antagonistic potential against phytopathogens through the release of active secondary metabolites and the induction of systemic resistance in plants by producing considerable amounts of surfactins (25).

It should be noted that the plant growth-promoting effects of these PGPR are influenced by environmental factors. Previous findings have shown that soil moisture, temperature, and the availability of an energy source (carbon) determine the activity of these organisms and soil microflora, which are important for the incidence of disease and suppression thereof (26). These data explained why only a limited number of strains have been successfully applied in agriculture and developed commercially as biofertilizers or biocontrol agents, although several *Bacillus* strains have been developed as biocontrol agents in for plant protection and improving plant growth (27). Thus, it is a substantial advantage that indigenous microorganisms have been developed as biocontrol agents and applied in local arable land.

In conclusion, we successfully selected an effective antagonistic bacterial strain (XH-9) from wheat rhizosphere and identified it as *B. amyloliquefaciens subsp. Plantarum*, based on phylogenetic analysis of the 16S rRNA sequence. Data from pot experiments indicated that the XH-9 strain increased in the growth of wheat and reduced the number of *F. oxysporum* in the treatment group. Antagonistic mechanisms and PGP characteristics were revealed in terms of nitrogen fixation, phosphate and potassium solubilization, and production of growth hormones, ACC deaminase, diffusible and volatile antibiotics, siderophores, cellulase, glucanase, protease, and chitinase. The key point is that the XH-9 strain can colonize the root system of different plant crops, which has been considered a relevant trait for biocontrol (28). Additionally, Chowdhury et al. (24) reported that plant growth-promoting activity is linked with the ability to suppress soil-borne plant pathogens. To our knowledge, bacteria exhibiting all known plant growth-promoting features simultaneously are rare (29). Fortunately, the XH-9 strain possesses all plant growth-promoting characteristics. Although further experiments are needed to determine the effectiveness of the XH-9 strain under field conditions, we have confidence that the XH-9 strain can be developed as a biocontrol agent for improving crop productivity in traditional and organic production systems.

## Experimental Procedures

### Soil sampling

Soil samples from healthy wheat plants were considered good sources of PGPR and were collected randomly from a wheat field located in Taian City (Shandong Province, China). One whole wheat plant was carefully uprooted during the grain-filling stage in 2015, the aerial parts were chopped off, and the residual root portion was placed in a polythene bag, labeled and tied, and further placed in a box containing ice. The ice box was immediately transported to a laboratory and stored in a refrigerator at 4°C for further studies.

### Isolation of bacterial strains

A mixed rhizosphere soil sample was collected from the roots as described previously (6). Ten grams of soil were suspended in 90 ml of sterile distilled water and mixed on a table concentrator for 30 min. The soil samples were serially diluted (up to 10^6^-fold), plated on PDA plates, and incubated at 28°C ± 2°C for 2-3 d. Bacterial colonies growing on the plates were isolated according to their different visual characteristics and maintained on PDA slants at 4°C for further studies (6).

### Screening and identification of antagonistic bacteria

All isolates were screened for antagonistic activity against soil-borne plant pathogenic fungi by performing dual culture assays. The isolates included member of the *Fusarium pseudograminearum*, *F. oxysporum, Alternaria* (Nees), *Colletotrichum gloeosporioides*, *Bipolaris sorokiniana, Botryosphaeria ribis*, and *Rhizoctonia solani* taxa, which are common phytopathogens and often cause soil diseases in various plants (30–33). The centers of PDA plates were inoculated with pure cultures of pathogens (5-mm discs) and incubated for 24 h. next; the bacterial isolates were incubated on both sides into 9-cm diameter Petri dishes containing PDA. Three replicate plates were maintained for each isolate. All plates were incubated at 30°C in a regulated incubator for 72 h. The percent growth inhibition was calculated using the following formula: inhibition (%) = 1− (fungal growth/control growth) × 100% (34).

### Ribosomal sequencing and phylogeny of isolates

The single most effective antagonistic bacterial strain (XH-9) showing a wide broad spectrum against phytopathogens was selected for further characterization, based on sequence analysis of 16S rDNA gene (11). The bacterium was grown in Luria Bertani(LB)broth culture at 28°C ± 2°C for 24 h and was then centrifuged at 3480 × g for 2 min. DNA was extracted using the EasyPure Bacteria Genomic DNA Kit (TransGen Biotech, Beijing, China) according to the manufacturer’s instructions. The 16S rDNA gene sequences from bacterial genomic DNA samples were amplified using universal primers 27F and 1492R (35). PCR amplification was performed in 50-μL reactions containing 25 μL of 2× Taq PCR Master Mix (TransGen Biotech, Beijing, China), 2 μL of each primer (10 μM), 2 μL of template DNA (10 ng/μL), and 19 μL of ddH_2_O. PCR amplification was performed under the following conditions: initial denaturation at 94°C for 3 min; 35 cycles of 94°C for 1 min, 54°C for 1 min, and 72°C for 1 min; and a final extension at 72°C for 10 min. The obtained amplicons were sequenced by a commercial sequencing company (Sangon Biotech, Shanghai, China). The obtained gene sequences were analyzed and BLAST searched againstwith the GenBank database (https://blast.ncbi.nlm.nih.gov/Blast.cgi), and a phylogenetic tree was constructed using MEGA 5.1 software (36).

### Antagonistic mechanisms

*Bipolaris sorokiniana*, *F. oxysporum*, and *Rhizoctonia solani* were used to test the diffusible and volatile antibiotics, as described previously (37). Chitinase activity was determined using colloidal chitin as the substrate (38); Glucanase activity was determined using yeast glucans as the substrate(39); Cellulase activity was determined using sodium carboxymethyl cellulose as the substrate(40).The amount of increased reducing sugars in the reaction mixture was determined using the dinitrosalicylic acid solution (41). Protease activity was determined by measuring tyrosine released from casein in the reaction mixture (42). Siderophore production was detected by the universal chrome azurol S) assay. An orange halo around the colonies grown on CAS media was considered as positive for the production of siderophores (43). Each experiment was performed in triplicate, and the results are expressed as mean values and standard deviations.

### Growth-promoting characteristics of the isolates

All selected antagonistic isolates were characterized for differences in their traits. Qualitative determination of phosphate and potassium solubilization by strains that induced a clear zone around colonies on NBRIP growth medium and silicate solid medium,which were considered as positive results for phosphate and potassium solubilization (44). Quantitative analysis of the phosphate- and potassium-solubilization efficiencies was conducted as described previously (45, 46). Nitrogenase activity was measured by performing a previously described(47).Salkowski assays were used to analyze IAA production in liquid cultures supplemented with L-tryptophan (0.5mg ml^−1^) for 48 h (48). Qualitative detection of ACC deaminase produced by the isolates was performed using DF-ACC salts minimal medium (49)

### Root-colonization assay

To examine the root-colonizing capacity of selected strains in vitro, the original XH-9 strain was replaced by a rifampicin- and spectinomycin-resistant mutant of the XH-9 strain, which was designated XH-9M. The process of introducing mutations was performed as described previously (50). The wheat seed surface was sterilized with 75% ethanol for 1 min, exposed to 1% sodium hypochlorite for 30 min, and rinsed thoroughly with sterile distilled water. Surface-sterilized wheat seeds were sown in 10-L plastic pots containing soil collected from which the XH-9 strain was isolated. The experiment involved a completely randomized block design with three replicates for each treatment, with 5 pots per replicate; each pot has 5 wheat seeds. The XH-9M strain was grown overnight in LB broth at 30°C and 200 rpm, after which the bacterial suspension was centrifuged at 1073 × *g* for 10 min and resuspended in phosphate-buffered saline (1 × 10^8^ CFUs/mL) prior to use. Twenty milliliters of bacterial suspension was poured into pots containing wheat seedlings that were germinated a week before (51). Bacteria were recovered from the rhizosphere by collecting wheat rhizosphere soil every 10 days after bacterial inoculation. Five-gram soil samples were placed into 250-mL conical flasks containing 50 mL of sterile water and mixed on a table concentrator for 30 min. The suspensions were serially diluted and plated on PDA agar containing rifampicin (300 μg mL^−1^) and spectinomycin (300 μg mL^−1^). The plates were incubated for 2–3 d at 30°C, the number of colonies was counted, and the total population was expressed as the CFU g^−1^ of dry soil. Furthermore, the colonization ability of the selected strains was also tested on corn and chili crops in the same manner.

### Pot experiment design and qPCR

In this study, qPCR was used to quantitatively measure the *F. oxysporum* biomass in wheat rhizosphere soil. Wheat seeds were sterilized and sown in clay tile pots, each of which had been inoculated with a 20-mL suspension of *F. oxysporum* conidia (10^7^ mL^−1^). After one week, the roots of plantlets were inoculated with 20 mL of bacterial suspension; the roots of the control plantlets were inoculated with the same volume of sterile water. After inoculation, both the treated and control rhizosphere soil were collected every 10 d and stored at −80°C to detect the amount of *F. oxysporum* by qPCR. The pot experiments followed a completely randomized block design with three replicates for each treatment, with 5 pots per replicate. Each pot had 5 wheat seeds. After 50 d, the wheat seedlings were uprooted separately to determine the biomass indices of the plants.

### Fungal and soil DNA extraction

Mycelia of fung collected from cultures grown on PDA medium were freeze-dried iquid nitrogen and ground to a fine powder using an autoclaved mortar and pestle (52). Next, DNA was extracted from 100-200 mg of the powdered mycelia using the HP Fungal DNA Kit (TransGen Biotech, Beijing, China).

### Primer selection *and specificity in vitro*

qPCR assays of *F. oxysporum* DNA were conducted using the following specific primer pair: JB-F (5’-CATACCACTTGTTGTCTCGGC-3’) and JB-R (5’-GAACGCGAATTAACGCGAGTC-3’), which was provided by Professor Zhiquan Mao of the College of Horticulture at Shandong Agricultural University. To evaluate the specificity of the primer pairs, genomic DNA samples were extracted from *F. pseudograminearum*, *F. graminearum, B. sorokiniana*, and *R. solani* and used as templates for PCR amplification, with sterile water used as a negative control. Each reaction mixture contained 25 μL of 2×Easy Taq PCR Super Mix (TransGen Biotech, Beijing, China), 2.0 μL of each primer, 2.0 μL of DNA template (10 ng/μL), and 19 μL of sterile distilled water. The thermal cycling conditions were as follows: initial denaturation at 95°C for 5 min, followed by 55 cycles of 95°C for 1 min and 54°C for 1 min, and a final extension at 72°C for 10 min. The PCR products were verified by running them on 2% agarose gels and visualizing them under UV light.

### Plasmid construction and extraction

Specific target fragment of *F. oxysporum* were amplified using the specific PCR primer pair, JB-F/JB-F. The PCR amplification system and thermo cycling program are described above. The amplicons were extracted from 1% agarose gels using the EasyPure Quick Gel Extraction Kit (TransGen Biotech, Beijing, China) and spectrophotometrically quantified in a BioPhotometer to determine the amplicon concentrations. Furthermore, these amplicons were used to build carriers by ligation into the pEASY-T1 cloning vector at a 7: 1 molar ratio and then transformed into phage-resistant, chemically competent Trans1-T1 cells. Moreover, these transformants were incubated for 1 h in LB medium at 37°C 200 rpm and plated on PDA medium containing 8 μL IPTG and 40 μL X-gal. The plates were incubated for 24 h at 30°C; several white colonies were selected and sequenced in a commercial sequencing company (Sangon Biotech, Shanghai, China).

### Real-time qPCR experiments

qPCR was performed with a BioRad CFX Manager real-time PCR machine using TIAN GEN SuperReal PreMix plus SYBR Green. Plasmids were serially diluted (up to 10^−6^-fold) and used as DNA standards, with each standard measured in triplicate independent experiments. Each amplification was carried out in a 15-μL volume containing 7.5 μL of 2× TransStart Tip Green qPCR SuperMix (TransGen Biotech, Beijing, China), 0.6 μL of each primer (10 μM), 1 μL of template DNA (10 ng/μL), and ddH_2_O added to 15 μL. A two step amplification program was performed as follows: initial denaturation at 94°C for 30 s, followed by 45 cycles of 94°C for 5 s, 60°C for 15 s, and 72°C for 10 s. Melting curve was made at 0.6°C temperature increments every 10 s within a temperature range of 60-100°C and carried out at 95°C for 10 s, 65°C for 5 s, and 95°C for 5 s. Ct values are were determined as the cycle number at which the PCR product was first fluorescently detected and was inversely related to log of the initial concentration, such that lower Ct values corresponded to higher initial DNA concentrations. A standard curve was developed by plotting the logarithm of known concentrations against the Ct values. To quantif the amount of *F. oxysporum* in the wheat rhizosphere soil, total soil DNA was extracted from wheat rhizosphere soil (treatment and control groups) every 10 d after bacterial inoculation and used as a DNA template for real-time qPCR. For each sample, 500 mg of rhizosphere soil was used to extract total soil DNA with a Soil DNA Kit (O Bio-Tek, Guangzhou, China). The PCR amplification system and thermocycling program were the same as described above, with each DNA sample being measured in triplicate independent experiments.

## Acknowledgements

This work was supported by the Special Fund for Agro-scientific Research in the Public Interest (no. 201503112) and the Forestry Industry Research special funds for public welfare projects (no. 201304212). The work was also supported by the Funds of from the Shandong forestry science and technology innovation project and Shandong “Double Tops” Program.

## Compliance with Ethical Standards

This article does not contain any studies with human participants or animals performed by any of the authors.

## Conflict of Interest

The authors have no conflict of interest to declare.

## References

1 Nourozian J, Etebarian HR, Khodakaramian, G. 2006. Biological control of Fusarium graminearum on wheat by antagonistic bacteria. SJST 28:29–38.

2 Li, E, Ling, J, Wang, G, Xiao, J, Yang, Y, Mao, Z, Wang, X, Xie, B. 2015. Comparative Proteomics Analyses of Two Races of Fusarium oxysporum f. sp. conglutinans that Differ in Pathogenicity. SCI REP-UK 5:13663.

3 Shanmugam, V, Kanoujia N. 2011. Biological management of vascular wilt of tomato caused by Fusarium oxysporum f.sp. lycospersici by plant growth-promoting rhizobacterial mixture. Biol Control 57:85–93.

4 Dufour, MC, Fontaine, S, Montarry, J, Corio-Costet MF. 2011. Assessment of fungicide resistance and pathogen diversity in Erysiphe necator using quantitative real-time PCR assays. Pest Manag Sci 67:60–69.

5 Jetiyanon, K, Kloepper, JW. 2002. Mixtures of plant growth-promoting rhizobacteria for induction of systemic resistance against multiple plant diseases. Biol Control 24:285–291.

6 Gopalakrishnan, S, Humayun, P, Kiran, BK, Kannan, IG, Vidya, MS, Deepthi, K, Rupela, O. 2011Evaluation of bacteria isolated from rice rhizosphere for biological control of charcoal rot of sorghum caused by Macrophomina phaseolina (Tassi) Goid. World J Microb Biot 27:1313–1321.

7 Kloepper, JW, Beauchamp, CJ. 1992. A review of issues related to measuring colonization of plant roots by bacteria. Can J Microbio 38:1219–1232.

8 Ajit, NS, Verma, R, Shanmugam, V. 2006. Extracellular chitinases of fluorescent pseudomonads antifungal to Fusarium oxysporum f. sp. dianthi causing carnation wilt. Curr Microbiol 52:310–316.

9 Kamilova, F, Leveau, JH, Lugtenberg, B. 2007. Collimonas fungivorans, an unpredicted in vitro but efficient in vivo biocontrol agent for the suppression of tomato foot and root rot. Environ Microbiol 9:1597–1603.

10 Schippers, B, Bakker, AW, Bakker, PAHM. 1987. Interactions of Deleterious and Beneficial Rhizosphere Microorganisms and the Effect of Cropping Practices. Annu Rev Phytopathol 25:339–358.

11 Khabbaz, SE, Abbasi, PA. 2014. Isolation, characterization, and formulation of antagonistic bacteria for the management of seedlings damping-off and root rot disease of cucumber. Can J Microbio 60:25–33.

12 Someya, N, Nakajima, M, Watanabe, K, Hibi, T, Akutsu, K. 2003. Influence of bacteria isolated from rice plants and rhizospheres on antibiotic production by the antagonistic bacterium Serratia marcescens strain B2. J Plant Pathol 69:342–347.

13 Minerdi, D, Bossi, S, Gullino, ML, Garibaldi, A. 2009. Volatile organic compounds: a potential direct long-distance mechanism for antagonistic action of Fusarium oxysporum strain MSA 35. Environ Microbiol 11:844–854.

14 Zhan-Bin, Wu, Lei-Fei, Liang, Chen, Feng, Yong-Jun, Yuan, Xing-Fang. 2013. Isolation, screening and identification of antagonistic downy mildew;endophytic bacteria from cucumber. Eur J Plant Pathol 137:847–857.

15 Cook, RJ. 2003. Making Greater Use of Introduced Microorganisms for Biological Control of Plant Pathogens. Annu Rev Phytopathol 31:53.

16 Kamilova, F, Leveau, JH, Lugtenberg, B. 2010. Collimonas fungivorans, an unpredicted in vitro but efficient in vivo biocontrol agent for the suppression of tomato foot and root rot. Environ Microbiol 9:1597–1603.

17 Camiloa, R, Josephw, K. 2010. Plant growth promotion by Bacillus amyloliquefaciens FZB45 depends on inoculum rate and P-related soil properties. Biol Fert Soils 46:835–844.

18 Harris, D, Breese, WA, Rao, JVDKK. 2005. The improvement of crop yield in marginal environments using ‘on-farm’ seed priming: nodulation, nitrogen fixation, and disease resistance. Aust J Agr Res 56:1211–1218.

19 Choi, O, Kim, J, Ryu, CM, Chang, SP. 2004. Colonization and Population Changes of a Biocontrol Agent, Paenibacillus polymyxa E681, in Seeds and Roots. Plant Pathology J 20:712–714 Vol. 712.

20 Qiao, JQ, Wu, HJ, Huo, R, Gao, XW, Borriss, R. 2014. Stimulation of plant growth and biocontrol by Bacillus amyloliquefaciens subsp. plantarum FZB42 engineered for improved action. Chemical & Biological Technologies in Agriculture 1:12.

21 Kang, SM, Radhakrishnan, R, Lee, IJ. 2015. Bacillus amyloliquefaciens subsp. plantarum GR53, a potent biocontrol agent resists Rhizoctonia disease on Chinese cabbage through hormonal and antioxidants regulation. World J Microb Biot 31:1517–1527.

22 Borriss, R. 2011. Use of Plant-Associated Bacillus Strains as Biofertilizers and Biocontrol Agents in Agriculture. Bacteria in Agrobiology Plant Growth Responses:41–76.

23 Szilagyi-Zecchin VJ, Mógor, ÁF, Ruaro, L, Röder, C. 2015. Tomato seedlings growth (Solanum lycopersicum) promoted by bacteria Bacillus amyloliquefaciens subsp. plantarum FZB42 in organic system. Revista De Ciencias Agrarias 38:26.

24 Chowdhury, SP, Hartmann, A, Gao, X, Borriss, R. 2015. Biocontrol mechanism by root associated Bacillus amyloliquefaciens FZB42 - a review. Front Microbiol 6:780.

25 Molinatto, G, Puopolo, G, Sonego, P, Moretto, M, Engelen, K, Viti, C, Ongena, M, Pertot, I. 2016. Complete genome sequence of Bacillus amyloliquefaciens subsp. plantarum S499, a rhizobacterium that triggers plant defences and inhibits fungal phytopathogens. J. Biotechnol. 238:56–59.

26 Gupta, VVSR, Rovira, AD, Roget, DK. 2011. Principles and Management of Soil Biological Factors for Sustainable Rainfed Farming Systems. Rainfed Farming Systems:149–184.

27 Solanki, MK, Kumar, S, Pandey, AK, Srivastava, S, Singh, RK, Kashyap, PL, Srivastava, AK, Arora, DK. 2012. Diversity and antagonistic potential of Bacillus spp. associated to the rhizosphere of tomato for the management of Rhizoctonia solani. Biocontrol Sci Techn 22:203–217.

28 Kamilova, F, Leveau, JH, Lugtenberg, B. 2007. Collimonas fungivorans, an unpredicted in vitro but efficient in vivo biocontrol agent for the suppression of tomato foot and root rot. Environ Microbiol 9:1597–1603.

29 Muthukumarasamy, R, Revathi, G, Seshadri, S, Lakshminarasimhan, C. 2002. G. diazotrophicus (syn. Acetobacter diazotrophicus), a promising diazotrophic endophyte in tropics. Curr Sci India 83:137–145.

30 Wildermuth, GB, Mcnamara, RB. 1991. Effect of cropping history on soil populations of Bipolaris sorokiniana and common root rot of wheat. Aust J Agr Res 42:779–790.

31 Vakalounakis, DJ. 1996. Root and stem rot of cucumber caused by Fusarium oxysporum f.sp. radicis-cucumerinum f.sp. nov. Plant Dis 80:313.

32 Obanor, F, Neate, S, Simpfendorfer, S, Sabburg, R, Wilson, P, Chakraborty, S. 2013. Fusarium graminearum and Fusarium pseudograminearum caused the 2010 head blight epidemics in Australia. Plant Pathol 62:79–91.

33 Wang, QH, Niu, SG, Liu, XH, Liu, YS. 2010. Identification of species bf-02 and its antifungal activity against several plant pathogens. Journal of Shandong Agricultural University 41:513–516.

34 Hameeda, B, Harini, G, Rupela, OP, Kumar Rao JV, Reddy, G. 2010. Biological Control of Chickpea Collar Rot by Co-inoculation of Antagonistic Bacteria and Compatible Rhizobia. Indian J. Microbiol. 50:419.

35 Watanabe, K, Kodama, Y, Harayama, S. 2001. Design and evaluation of PCR primers to amplify bacterial 16S ribosomal DNA fragments used for community fingerprinting. J Microbiol Meth 44:253–262.

36 Mendez, J. 2014. Characterization of phosphate-solubilizing bacteria isolated from the arid soils of a semi-desert region of north-east Mexico. Biol Agric Hortic 30:211–217.

37 Asha, BB, Nayaka, SC, Shankar, ACU, Srinivas, C, Niranjana, SR. 2011. Selection of effective bio-antagonistic bacteria for biological control of tomato wilt caused by fusarium oxysporum f. sp. lycopersici. Bioscan An International Quarterly Journal of Life Sciences.

38 Jankiewicz, U, Brzezinska, MS, Saks, E. 2012. Identification and characterization of a chitinase of Stenotrophomonas maltophilia, a bacterium that is antagonistic towards fungal phytopathogens. J Biosci Bioeng 113:30–35.

39 Pérez, LM, Besoaín X, Reyes, M, Pardo, G, Montealegre, J. 2002. The expression of extracellular fungal cell wall hydrolytic enzymes in different Trichoderma harzianum isolates correlates with their ability to control Pyrenochaeta lycopersici. Biol Res. 35:401.

40 Miller, GL. 1959. Use of Dinitrosalicylic Acid Reagent for Determination of Reducing Sugar. Anal Biochem 31:426–428.

41 Horikoshi, K. 1973. Comparative studies on beta-1,3-glucanases of microorganisms. Yeast Mould An Plant Protoplasts Proceedings.

42 Bian HH, Huang, Wang. 2009. Protease activity and chyme protein tracing by GFP in the digestive tract of flounder Paralichthys olivaceus. Progress in Fishery Sciences.

43 Schwyn, B, Neilands, JB. 1987. Universal chemical assay for the detection and determination of siderophores. Anal Biochem 160:47–56.

44 Mehta, S, Nautiyal, CS. 2001. An efficient method for qualitative screening of phosphate solubilizing bacteria. Curr Microbiol 43:51–56.

45 Bray, RH, Kurtz, LT. 1945. Determination of Total, Organic, and Available Forms of Phosphorus in Soils. Soil Sci 59:39–46.

46 LüChengQun, Huang, BL. 2010. Isolation and characterization of azotobacteria from pine rhizosphere. Afr J Microbiol Res 4:1299–1306.

47 Marra, LM, Oliveira, SMD, Ferreira, PAA, Soares, BL, Carvalho, RDF, Lima, JMD, Moreira, FMDS. 2012. Biological nitrogen fixation and phosphate solubilization by bacteria isolated from tropical soils. Plant Soil 357:289–307.

48 Abiala, MA, Odebode, AC, Hsu, SF, Blackwood, CB. 2015. Phytobeneficial Properties of Bacteria Isolated from the Rhizosphere of Maize in Southwestern Nigerian Soils. Appl Environ Microb 81:4736–4743.

49 Husen, E, Wahyudi, AT, Suwanto, A, Saraswati, R. 2009. Soybean seedling root growth promotion by 1-aminocyclopropane-1-carboxylate deaminase-producing pseudomonads. Indonesian Journal of Agricultural Science 10:19–25.

50 Ji, X, Lu, G, Gai, Y, Gao, H, Lu, B, Kong, L, Mu, Z. 2010. Colonization of Morus alba L. by the plant-growth-promoting and antagonistic bacterium Burkholderia cepacia strain Lu10-1. BMC Microbiol, 10,1(2010-09-20) 10:1–12.

51 Someya, N, Nakajima, M, Akutsu, K. 2010. Potential of an Antagonistic Bacterium Serratia marcescens Strain B2 for the Biological Control of Cucumber Damping-Off Disease. Biocontrol Sci 10:101–104.

52 Perezartes, E, Garciapedrajas, MD, Bejaranoalcazar, J, Jimenezdiaz, RM. 2000. Differentiation of cotton-defoliating and nondefoliating pathotypes of Verticillium dahliae by RAPD and specific PCR analyses. Eur J Plant Pathol 106:507–517.

